# Platelet TSP-1 Controls Prostate Cancer-Induced Osteoclast Differentiation and Bone Marrow-Derived Cell Mobilization through TGFβ-1

**DOI:** 10.1101/2020.02.11.943860

**Authors:** Bethany A. Kerr, Koran S. Harris, Lihong Shi, Jeffrey S. Willey, David R. Soto-Pantoja, Tatiana V. Byzova

**Author notes:** Corresponding Author: Bethany Kerr, Ph.D., Wake Forest School of Medicine, Medical Center Blvd, Winston-Salem, NC, 27157.; Twitter: @BethanyKerrLab.

## Abstract

The development of distant metastasis is the main cause of prostate cancer (CaP)-related death with the skeleton being the primary site of metastasis. While the progression of primary tumors and the growth of bone metastatic tumors are well described, the mechanisms controlling pre-metastatic niche formation and homing of CaP to bone remain unclear. Through prior studies, we demonstrated that platelet secretion was required for ongoing tumor growth and pre-metastatic tumor-induce bone formation and bone marrow-derived cell mobilization to cancers supporting angiogenesis. We hypothesized that proteins released by the platelet α granules were responsible for inducing changes in the pre-metastatic bone niche. We found that the classically anti-angiogenic protein thrombospondin (TSP)-1 was significantly increased in the platelets of mice bearing tumors. To determine the role of increased TSP-1, we implanted tumors in TSP-1 null animals and assessed changes in tumor growth and pre-metastatic niche formation. TSP-1 loss resulted in increased tumor size and enhanced angiogenesis but reduced bone marrow-derived cell mobilization and tumor-induced bone formation with enhanced osteoclast formation. We hypothesized that these changes in the pre-metastatic niche were due to the retention of TGF-β1 in the platelets of mice with TSP-1 deleted. To assess the importance of platelet-derived TGF-β1, we implanted CaP tumors in mice with platelet-specific deletion of TGF-β1. Similar to TSP-1 deletion, loss of platelet TGF-β1 resulted in increased angiogenesis with a milder effect on tumor size and BMDC release. Within the bone microenvironment, platelet TGF-β1 deletion prevented tumor-induced bone formation due to increased osteoclastogenesis. Thus, we demonstrate that the TSP-1/TGF-β1 axis regulates pre-metastatic niche formation and tumor-induced bone turnover. Targeting the platelet release of TSP-1 or TGF-β1 represents a potential method to interfere with the process of CaP metastasis to bone.

## INTRODUCTION

An estimated 191,930 men will be diagnosed with prostate cancer (CaP) in 2020, surpassing even lung cancer in incidence [1]. Early detection has improved survival rates to 100 percent with a local diagnosis. However, men diagnosed with distant CaP metastases face a 30 percent chance of survival at five years, with bone being the most common site of CaP metastasis [1]. In fact, 90% of CaP patients display skeletal metastases at autopsy regardless of prior bone symptom reporting. Bone metastases are responsible for severe bone pain, decreased mobility, fractures, spinal cord compression, and hypercalcemia resulting in patient morbidity. Metastatic skeletal CaP lesions can produce many growth factors and cytokines which alter the bone structure. Several studies have described alterations in both osteoclast and osteoblast activity in early metastasis, with increased bone formation prevailing in late metastasis [2, 3]. While the effects of CaP tumor growth in bone have been studied, the mechanisms of CaP metastasis to skeletal tissues remain unclear. We have shown that prior to metastasis, there is cross-talk between the tumor and bone, which may account for the predilection of CaP to metastasize to the skeleton [4–6]. Also, this communication indicates the preparation of a pre-metastatic niche for the circulating tumor cells.

Possible mechanisms for communication between the primary tumor and bone include proteins, RNAs, and mRNAs circulating in plasma, within platelets, or attached to sentinel tumor cells. Platelets function in metastasis by aggregating with tumor cells, coating circulating tumor cells as protection from immune recognition, promoting tumor cell adhesion and growth, enhancing angiogenesis, and stabilizing tumor vasculature [7–11]. In the primary tumor, platelets promote tumor stability and metastasis by direct interaction with tumor cells, with the tumor vasculature, and through secreted cytokines [12]. During tumor progression to metastasis, our studies have shown that platelets are required for increased bone formation stimulated by primary tumor growth [5] and also for mobilization of bone marrow-derived cells (BMDCs) from the bone marrow [6]. In the bone microenvironment, recent studies have demonstrated the ability of platelets to induce fracture healing and a dose-dependent effect on bone regeneration [13]. However, other studies have shown that factors released by platelet activation, such as matrix metalloproteinase (MMP)-9, cathepsin K, tartrate-resistant acid phosphatase (TRAP), receptor activator of NF-κB ligand (RANKL) and osteoprotegerin, may induce the formation of osteoclasts [14]. Importantly, platelets have selective separation, uptake, and release of pro-and anti-angiogenic proteins [15–17]. This controlled and specific uptake and release indicates that platelets may have an active role in tumor progression and the development of distant metastases.

Platelets contain many pro-angiogenic and anti-angiogenic factors: thrombospondin (TSP)-1, vascular endothelial growth factor (VEGF), transforming growth factor (TGF)-β1, in addition to the proteins capable of altering bone formation. TSP-1 controls in both bone turnover and CaP progression. In lung and breast cancers, TSP-1 promotes tumor cell metastasis, and adhesion [18, 19], with one of the studies demonstrating that cell metastasis due to TSP-1 is dependent upon platelets [18]. In breast cancer patients, TSP-1 levels in the plasma correlate directly with cancer progression and severity [20]. Functionally, TSP-1 binds latent TGF-β1 and induces its activation [21, 22], which may account for many of TSP-1’s effects on cancer growth and bone turnover. TGF-β1 levels in CaP patients are predictive of future biochemical recurrence [23–25] and are elevated in patients with bone metastases [24, 26]. Plasma TGF-β1 levels are largely due to the release of platelet TGF-β1 during the collection and preparation of blood samples [27] indicating that the TGF-β1 associated with CaP bone metastasis is likely platelet-derived. The utility of platelet TSP-1 and TGF-β1 as cancer biomarkers presents the possibility that these platelet proteins might play a role in pre-metastatic niche formation and tumor-induced bone formation.

While TSP-1 and TGF-β1 play important roles in the growth of primary tumors and bone metastases, the role of the TSP-1/TGF-β1 axis in pre-metastatic tumor communication remains unstudied. In this study, we investigated whether TSP-1 accumulation in platelets affects primary tumor progression and tumor-induced bone formation. We then examined what role the TSP-1 induced activation of TGF-β1 played in these pre-metastatic processes. We demonstrate that the platelet TSP-1/TGF-β1 axis has little effect on primary tumor growth but controls the pre-metastatic changes in the bone microenvironment. Thus, targeting TSP-1 and/or TGF-β1 will have the greatest effect in preventing bone metastasis rather than in inhibiting primary CaP growth.

## RESULTS

### Platelets accumulate TSP-1 during prostate tumor growth

We have previously shown that platelets actively sequester tumor-derived proteins involved promoting angiogenesis and bone marrow-derived cell (BMDC) mobilization [4, 6]. Using a syngeneic tumor model, we determined the levels of thrombospondin (TSP)-1 in platelets of mice bearing RM1 prostate tumors (Figure 1A). While TSP-1 levels remained unchanged in plasma, TSP-1 was approximately 6-fold higher in the platelets of mice bearing tumors compared with controls (Figure 1B). A similar upregulation of platelet TSP-1 was found in B16F10 syngeneic melanoma tumor growth [28]. Since TSP-1 is classically anti-angiogenic, we questioned whether this upregulation was solely a host response inhibiting tumor progression or if there was a secondary purpose of this significant upregulation.

**Figure 1.**
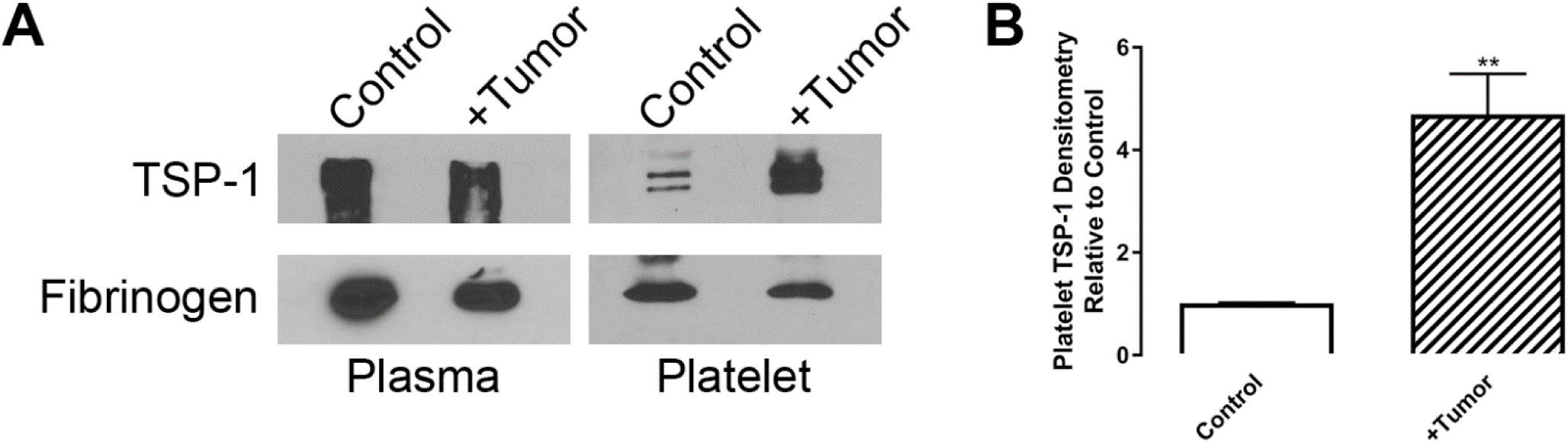
Tumor-bearing mice accumulate platelet TSP-1. (A-B) Activated platelets and plasma from control and RM1 tumor-bearing WT mice were immunoblotted for TSP-1 and fibrinogen. Densitometry of platelet TSP-1 bands was quantified normalized to loading controls and represented as mean relative to control±SEM (*n*=3).

### TSP-1 inhibits primary tumor growth but stimulates the pre-metastatic bone response

To determine how TSP-1 accumulation affects tumor progression, we implanted murine RM1 prostate tumors in TSP-1 null mice. Even in the presence of tumors, no TSP-1 was found in the platelets of TSP-1 null mice (Supplementary Figure 1). Consistent with previous studies, TSP-1 deletion resulted in significantly larger tumors compared with WT mice (Figure 2A). Analysis of cellular blood components demonstrated that TSP-1 deletion results in increased numbers of circulating lymphocytes and platelets (Supplementary Figure 2) compared with WT mice, consistent with previous studies [29]. Interestingly, the numbers of lymphocytes and platelets were decreased in mice bearing tumors (Supplementary Figure 2). In previous studies, we have shown that tumor progression is associated with pre-metastatic changes in the bone microenvironment resulting in the release of CXCR4^+^ bone marrow-derived cells (BMDCs) and bone formation [5, 6]. Also, we previously demonstrated the presence of circulating CD117^+^ cells in the blood is indicative of cancer progression in CaP patients [30]. Similar to our previous studies, RM1 tumor implantation in WT mice resulted in increased numbers of circulating CXCR4^+^ and CD117^+^ cells (Figure 2B). However, in this study, although tumors from TSP-1 null mice were significantly larger, we found decreased numbers of CXCR4^+^ and CD117^+^ cell numbers in the bloodstream (Figure 2B). This paradox indicates that TSP-1 may be anti-tumorigenic locally but promoting pre-metastatic changes in the bone microenvironment.

**Figure 2.**
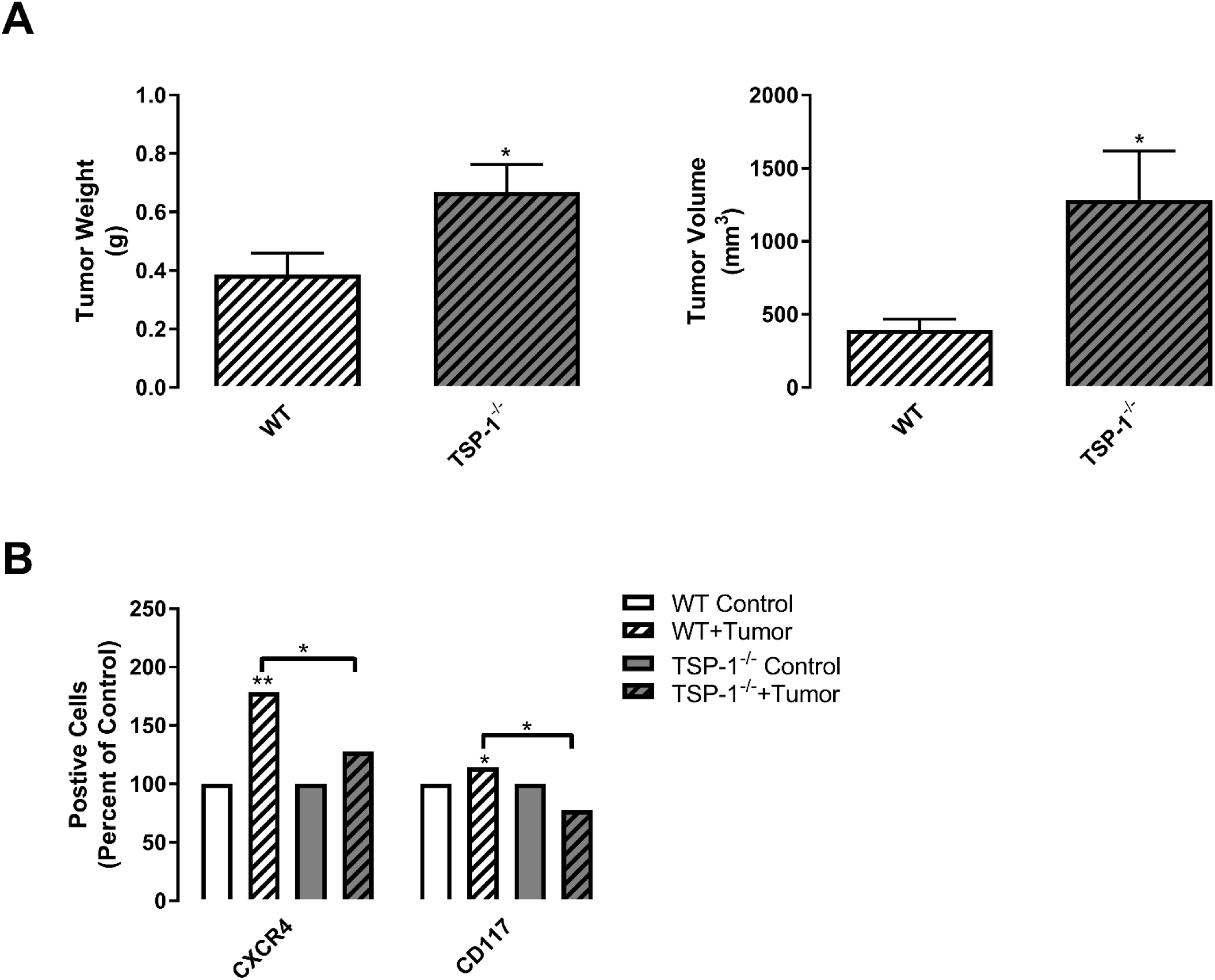
TSP-1 deficiency induces tumor formation but inhibits progenitor cell mobilization. WT and TSP-1 null mice were implanted with RM1 tumors subcutaneously. (A) Tumor weight and volume were measured and represented as mean±SEM (*n*=6-8). (B) Whole blood was collected and analyzed by flow cytometry for circulating progenitor cells using the markers CXCR4 and CD117. Quantification of positive cell numbers as mean percent of control±SEM (*n*=2-4). * represents p<0.05 and ** represents p<0.01 by Student’s *t* test (A) or one-way ANOVA (B).

To further examine the effect of TSP-1 deletion on the primary tumor, we assessed tumor sections for angiogenesis and platelet infiltration. Microvessel area was ∼1.3-fold higher in tumors in mice with global TSP-1 deletion (Figure 3A) corresponding to the loss of the anti-angiogenic protein. Analysis of mature smooth muscle actin (SMA)-positive blood vessels demonstrated ∼2.4-fold increased blood vessel infiltration in tumors from TSP-1 null mice compared with tumors from WT mice (Figure 3B). While circulating platelet number was decreased in mice bearing RM1 tumors, the numbers in TSP-1 null mice were not significantly different compared with WT mice (Supplementary Figure 2). However, within tumors, TSP-1 deletion resulted in ∼3.6-fold increased platelet numbers within the tumor. (Figure 3C.) Thus, the increased angiogenesis may permit more platelets to enter tumors, and TSP-1 expression might reduce platelet infiltration.

**Figure 3.**
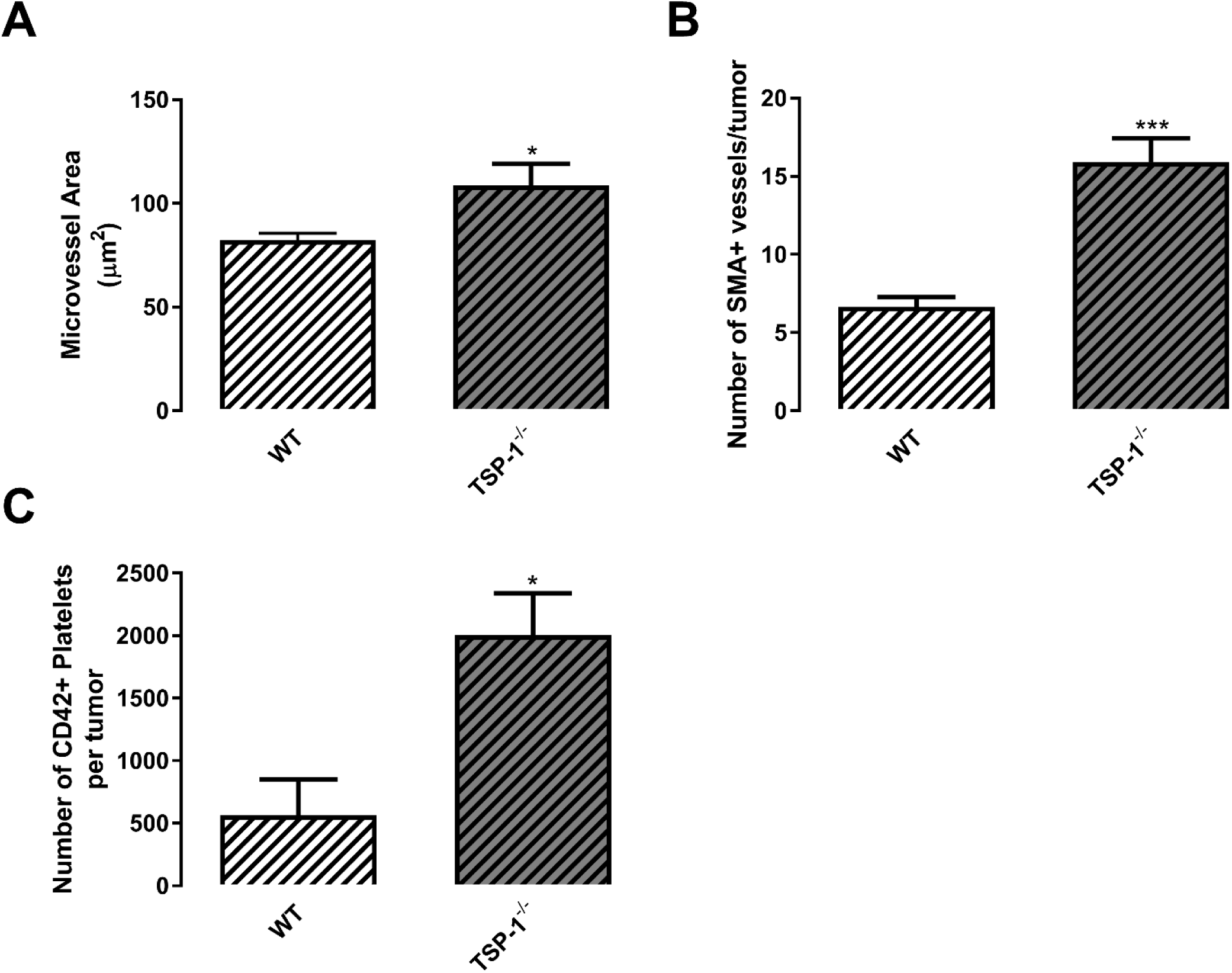
Loss of TSP-1 promotes tumor angiogenesis and platelet infiltration. (A-C) Tumors from WT and TSP-1 null mice were sectioned and stained with CD31 for microvessels, SMA for mature vessels, and CD42 for platelets. Quantification of staining per tumor is represented as mean±SEM (*n*=4). * represents p<0.05 and *** represents p<0.005 by Student’s *t* test.

To evaluate how TSP-1 regulates pre-metastatic niche formation, we analyzed changes in the bone structure by microCT. Pre-metastatic tumor progression stimulates bone formation in WT mice (Figure 4A), corresponding to previous studies [5]. However, in TSP-1 null mice, bone formation in response to tumor growth was decreased ∼1.5-fold (Figure 4A). TSP-1 null mice were shown to have mild chondrodysplasia with slightly disorganized columns. However, these changes do not affect the overall bone structure or growth [31]. Interestingly plasma calcium content (Figure 4B) and plasma alkaline phosphatase (Figure 4C), associated with bone formation, were not significantly different between WT and TSP-1 null mice bearing tumors indicating that osteoblast activity was equal in response to tumor growth. Conversely, platelet RANKL (Figure 4D) and MMP-9 (Figure 4E) were significantly increased in TSP-1 null mice compared with WT mice bearing tumors, indicating that osteoclast differentiation and function are significantly higher in TSP-1 null mice. Thus, TSP-1 may be required for the induction of bone formation in response to primary tumor growth.

**Figure 4.**
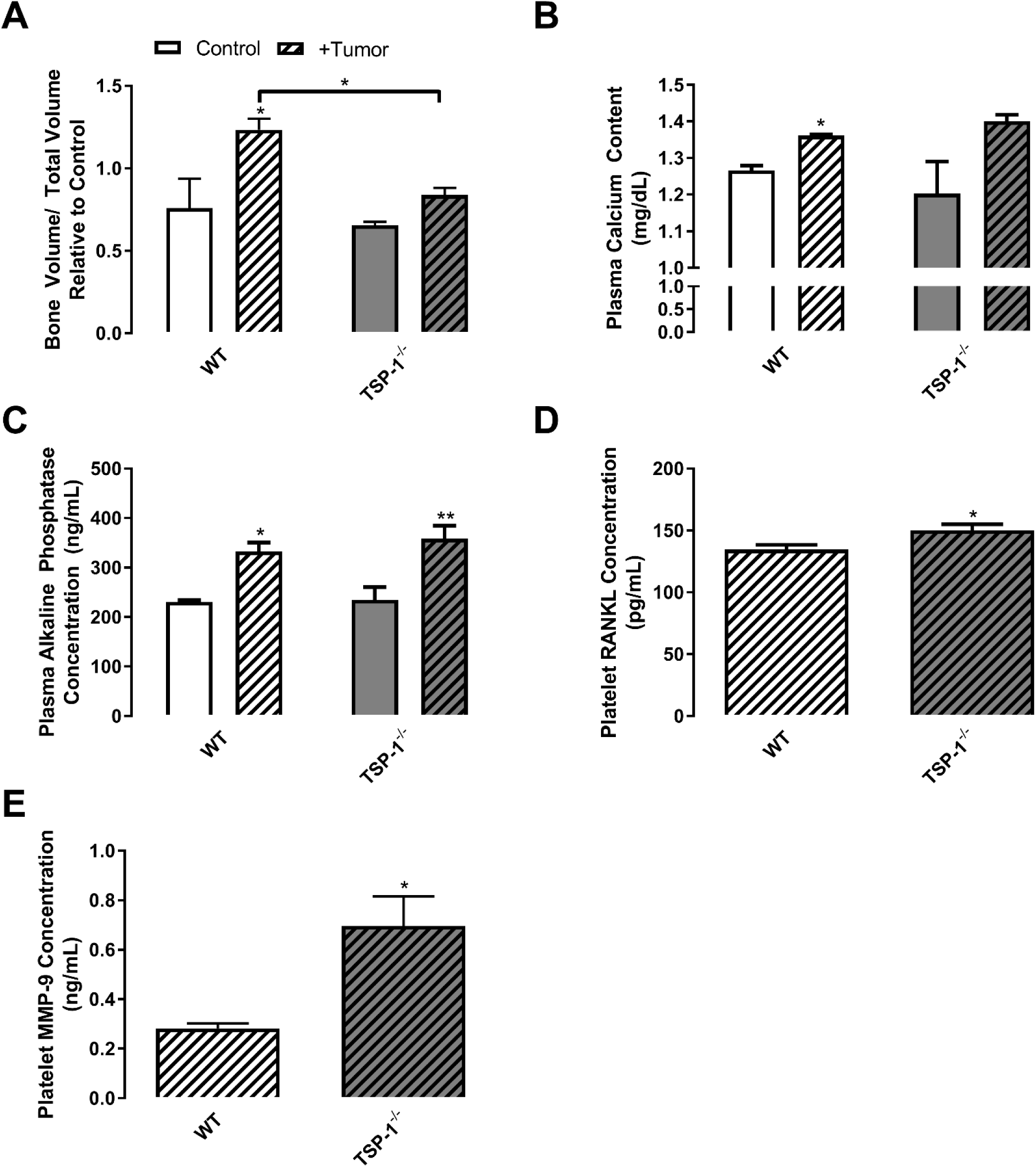
TSP-1 is required for tumor-induced bone formation. (A) Control and RM1 tumor-bearing WT and TSP-1 null mice were analyzed by microCT. The bone volume to total volume ratio was calculated and represented as mean±SEM (*n*=3-5). (B-E) Plasma and activated platelets were isolated and analyzed by ELISA for protein concentrations of calcium (B), alkaline phosphatase (C), RANKL (D), and MMP-9 (E) represented as mean±SEM (*n*=3-6). * represents p<0.05 and ** represents p<0.01 by one-way ANOVA (A-C) and Student’s *t* test (D-E).

### TSP-1 controls osteoclast differentiation and function in response to tumors

To further examine osteoclast differentiation after TSP-1 deletion, we sectioned bones of WT and TSP-1 null control and tumor-bearing mice and used TRAP staining to enumerate osteoclasts in the bone microenvironment. Quantification of osteoclast numbers demonstrated that TSP-1 null mice bearing tumors had 1.8-fold more osteoclasts on the bone surface compared with WT mice bearing tumors (Figure 5A). Also, bone marrow macrophages were isolated from control and tumor-bearing WT and TSP-1 null mice and differentiated *in vitro* into osteoclasts. The numbers of TRAP-positive cells with more than three nuclei were considered to be osteoclasts. Tumor-bearing mice bone marrow macrophages produced 3.7-and 1.9-fold more osteoclasts compared to control WT and TSP-1 null mice, respectively (Figure 5B). However, TSP-1 null control mice already contained 2.8-fold more osteoclast progenitors compared with WT (Figure 5B). Thus, the deletion of TSP-1 in the bone microenvironment results in increased osteoclast progenitor cells that can then be induced into further differentiation by tumor growth resulting in the prevention of the tumor-induced bone formation seen in WT mice.

**Figure 5.**
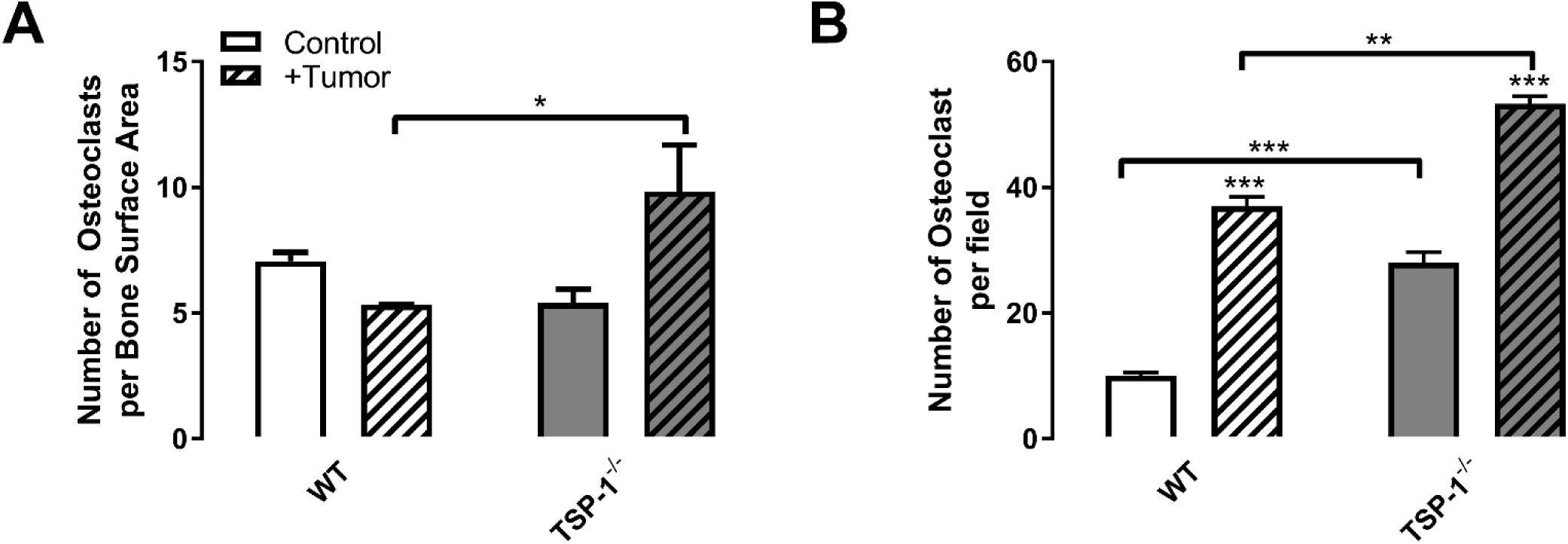
The loss of TSP-1 stimulates osteoclast differentiation. (A) Sections of bones isolated WT and TSP-1 null mice were stained for TRAP expression and the number of osteoclasts per bone surface area calculated and represented as mean±SEM (*n*=3). (B) BM macrophages isolated from control and RM1 tumor-bearing WT and TSP-1 null mice were differentiated towards osteoclasts by treatment with MCSF and RANKL. The number of osteoclasts per field was quantified as mean±SEM (*n*=3). * represents p<0.05, ** represents p<0.01 and *** represents p<0.005 by one-way ANOVA.

### Platelet TGF-β1 accumulated with tumor growth and TSP-1 deletion

TSP-1 binds latent TGF-β1 converts TGF-β1 to its active form [21, 22] in particular during the release of TSP-1 and TGF-β1 by alpha granules during platelet activation [32]. Accordingly, we examined the levels of latent TGF-β1 in platelets isolated from control and tumor-bearing WT and TSP-1 null mice (Figure 6A). Platelet TGF-β1 concentration was increased 2.2-fold in WT mice bearing tumors (Figure 6A), similar to increases seen in the platelets of advanced CaP patients (Supplementary Figure 3). As expected, TSP-1 deficiency resulted in the accumulation of TGF-β1 in platelets since without TSP-1, TGF-β1 could not be activated and released [22,32,33]. Correspondingly, TSP-1 null mice have reduced active TGF-β1 in their serum and platelets [34]. Platelet TGF-β1 levels are 5.0-fold higher in control TSP-1 null mice compared with WT (Figure 6A). Tumor-bearing TSP-1 null mice have 3.0-fold higher platelet TGF-β1 concentrations compared with their WT counterparts (Figure 6A).

**Figure 6.**
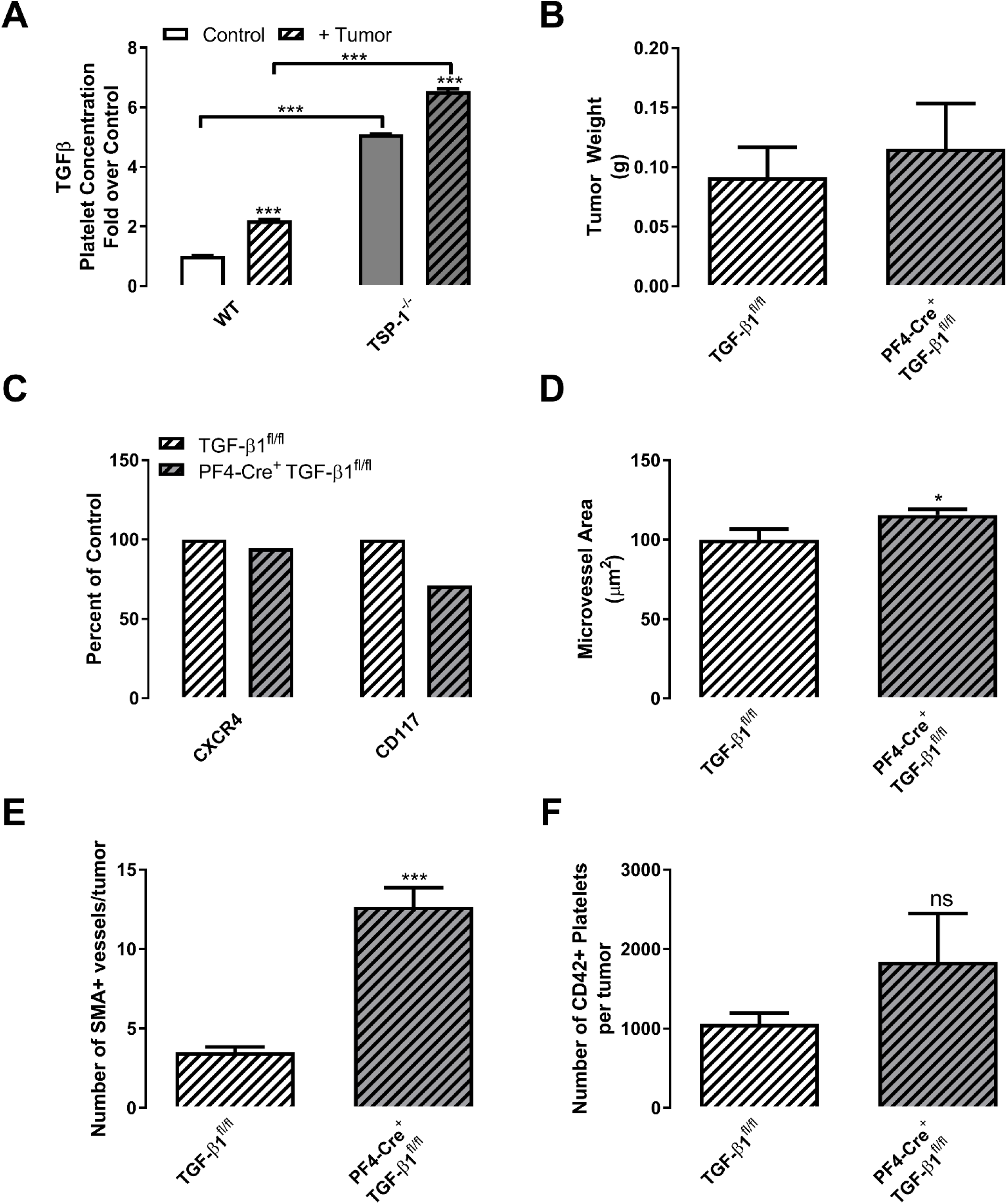
Platelet TGF-β1 is increased in TSP-1 null mice, and its deficiency inhibits progenitor cell mobilization and increases tumor angiogenesis. (A) Platelet TGF-β1 concentration was measured from control and RM1 tumor-bearing WT and TSP-1 null mice and represented as mean±SEM (*n*=3). (B) RM1 tumors from TGF-β1^fl/fl^ and PF4-Cre^+^;TGF-β1^fl/fl^ mice were weighed and represented as mean±SEM (*n*=3-11). (C) Whole blood was collected and analyzed by flow cytometry for circulating progenitor cells using the markers CXCR4 and CD117. Representative quantification of positive cell numbers as mean percent of control (*n*=2). (D) Tumors were sectioned and stained with CD31 for microvessels, SMA for mature vessels, and CD42 for platelets. Quantification of staining per tumor is represented as mean±SEM (*n*=4). * represents p<0.05, ** represents p<0.01, and *** represents p<0.005 by one-way ANOVA (A-B) or Student’s *t* test (D-F).

### Platelet TGF-β1 alters angiogenesis and tumor-induced bone formation without affecting primary tumor size

To determine the role of this increased platelet TGF-β1 in tumor progression, we generated megakaryocyte/platelet-specific TGF-β1 knockout mice (PF4-Cre^+^TGF-β1^fl/fl^) with littermate controls (TGF-β1^fl/fl^). PF4-Cre;TGF-β1^fl/fl^ mice have normal platelet counts and bleeding times [11]. The platelet/megakaryocyte specific deletion results in 93% decreased serum TGF-β1 levels and a 94% decrease in TGF-β1 release from platelets [27]. Tumors implanted in control TGF-β1^fl/fl^ and PF4-Cre^+^TGF-β1^fl/fl^ grew equally and reached similar volumes (Figure 6B). Elevated TGF-β1 levels in the circulation are correlated with increased numbers of circulating tumor cells in the bloodstream [35]. In concert, the deletion of TGF-β1 in platelets resulted in fewer numbers of circulating CXCR4^+^ and CD117^+^ progenitor cells (Figure 6C). Angiogenesis, as determined by microvascular area, was slightly increased in tumors from PF4-Cre^+^TGF-β1^fl/fl^ mice (Figure 6D). However, when tumor angiogenesis was examined by counting SMA^+^ mature blood vessels, PF4-Cre^+^TGF-β1^fl/fl^ tumor displayed 3.6-fold increased vasculature (Figure 6E), similar to that seen with TSP-1 deletion (Figure 3A). Platelet infiltration was not significantly altered with platelet-specific deletion of TGF-β1 (Figure 6F).

Although the deletion of platelet TGF-β1 did not significantly alter primary tumor size, we wanted to assess the effects of platelet-specific TGF-β1 loss on tumor-induced bone formation. As in WT mice, control TGF-β1^fl/fl^ mice displayed increased bone formation in response to tumor growth (Figure 7A). The deletion of platelet TGF-β1 prevented tumors of equal size (Figure 6B) from stimulating the same increase in bone (Figure 7A). This prevention of bone formation was similar to that seen with TSP-1 deletion. Thus, platelet TGF-β1 drives bone formation in response to tumors.

**Figure 7.**
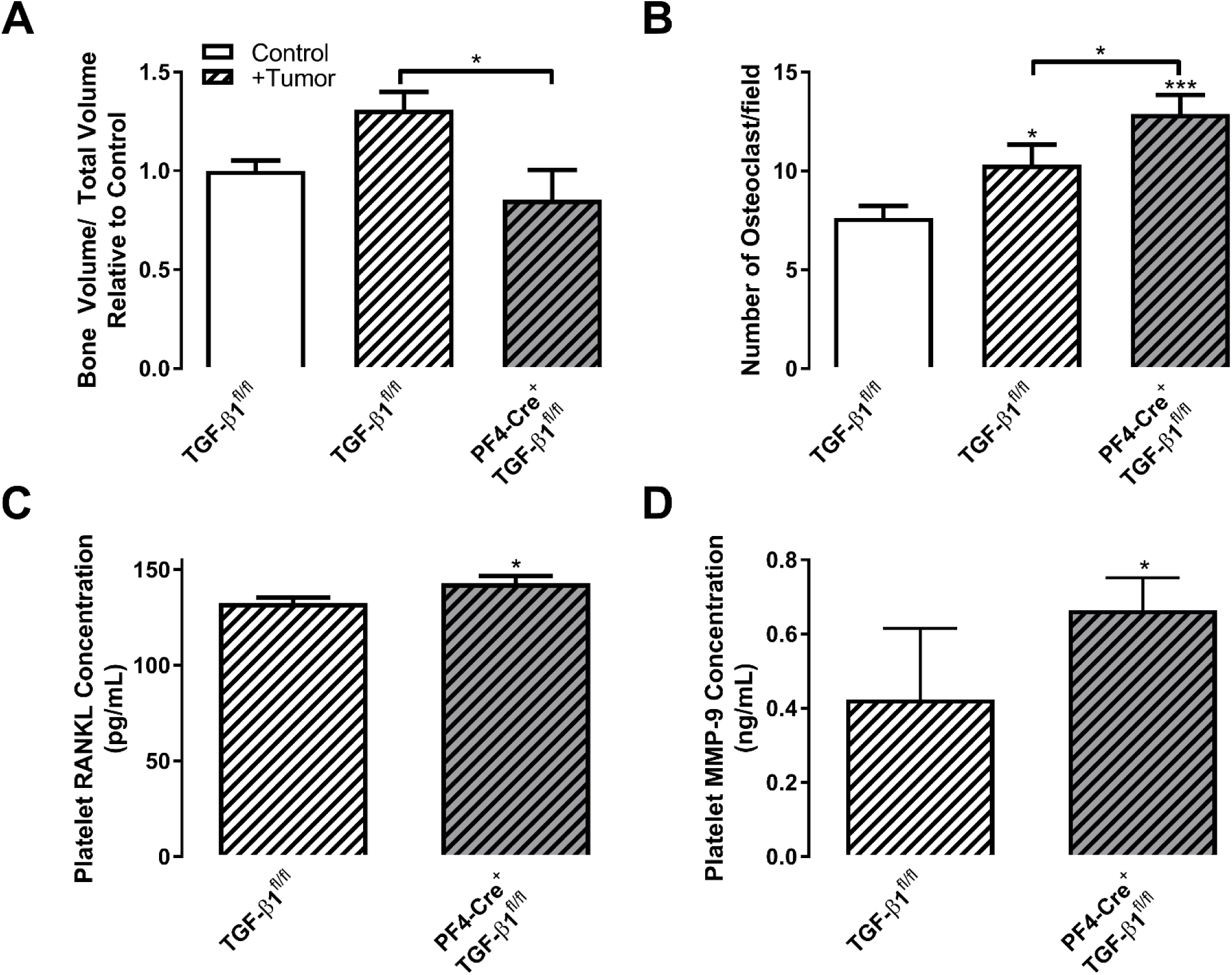
Platelet TGF-β1 is required for tumor-induced bone formation and osteoclast inhibition. (A) Control and RM1 tumor-bearing TGF-β1^fl/fl^ and PF4-Cre^+^;TGF-β1^fl/fl^ mice were analyzed by microCT. The bone volume to total volume ratio was calculated and represented as mean±SEM (*n*=3). (B) BM macrophages isolated from control and RM1 tumor-bearing TGF-β1^fl/fl^ and PF4-Cre^+^;TGF-β1^fl/fl^ mice were differentiated towards osteoclasts by treatment with MCSF and RANKL. The number of osteoclasts per field was quantified as mean±SEM (*n*=16). (C-D) Activated platelets were isolated and analyzed by ELISA for protein concentrations of RANKL (C) and MMP-9 (D) represented as mean±SEM (*n*=3-6). * represents p<0.05 and *** represents p<0.005 by one-way ANOVA (B) and Student’s *t* test (A, C-D).

Using the osteoclast differentiation assay from isolated bone marrow macrophages from mice, tumor growth in control TGF-β1^fl/fl^ mice resulted in enhanced numbers of osteoclast precursors (1.3-fold, Figure 7B). This effect was magnified another 1.2-fold in platelet TGF-β1 depleted mice (Figure 7B), similar to the effect seen with TSP-1 deletion. Correspondingly, the circulating platelet levels of RANKL, which controls osteoclast differentiation, were significantly higher in tumor-bearing mice with platelet TGF-β1 deletion compared with littermate controls (Figure 7C). In addition, levels of MMP-9, a marker for bone resorption and osteoclast recruitment [36], were increased 1.6-fold in platelet TGF-β1 depleted mice compared with controls (Figure 7D). Thus, platelet TGF-β1 is required for tumor-induced bone formation. Since TSP-1 controls TGF-β1 activation and release from platelets, the platelet TSP-1/TGF-β1 axis controls bone turnover and pre-metastatic niche formation in response to primary tumor growth.

## DISCUSSION

In this study, we examined the role of platelet TSP-1 in the preparation of the pre-metastatic niche during CaP progression. Loss of anti-angiogenic TSP-1 resulted in increased primary tumor growth and angiogenesis but prevented tumor communication with the bone microenvironment. Thus, there were decreases in circulating bone marrow-derived progenitor cells and tumor-induced bone formation despite an increase in primary tumor size. These effects were dependent upon TSP-1 activation of TGF-β1. Platelet-specific TGF-β1 deletion recapitulated TSP-1 loss, with diminished progenitor cell mobilization and tumor-induced bone formation. With the deletion of total TSP-1 or platelet-specific TGF-β1, osteoclast differentiation was enhanced tipping the balance away from the tumor-induced bone formation that would otherwise occur. Thus, targeting the TSP-1/TGF-β1 axis may be beneficial in preventing the metastatic progression of CaP.

The role of the TSP-1/TGF-β1 axis in cancer is debated in the literature with the source and tissue location of TSP-1 secretion altering its function. While our study does not evaluate the source of TSP-1 or TGF-β1 sequestered within platelets, another study demonstrated that increased TSP-1 in platelets is a result of stromal tissue upregulation as implantation of either TSP-1 expressing Lewis lung carcinomas or TSP-1 deficient B16F10 melanoma cells led to increased platelet TSP-1 [28]. This study further reported that TSP-1 was deposited into platelets by megakaryocytes in the bone marrow [28], implicating communication between primary tumors and the bone microenvironment similar to our study. For this reason, we utilized the platelet-factor4 Cre recombinase expressing mice to delete TGF-β1 in both megakaryocytes and platelets. However, the exact source of both TSP-1 and TGF-β1 would be better confirmed using xenograft models.

During development, prostates in TSP-1 null mice have dysregulated epithelial growth, with hyperplasia in the ventral and dorsal lobes. In the dorsal lobe of the murine prostate, considered to be the most analogous tissue to the human prostate peripheral zone where a majority of CaPs arise, TSP-1 deficiency results in a significant decrease in TGF-β1 activation. Further, castration induced TSP-1 expression in prostate epithelial cells in both humans and mice stimulating TGF-β1 activation [37]. In CaP, androgen withdrawal induces TSP-1 production in the prostate and while TSP-1 initially reduces tumor progression and vascularization, prolonged TSP-1 exposure results in high VEGF production by CaP cells and a dependency of the cancer cells on TSP-1 activated TGF-β1 [38, 39]. Thus, castration may stimulate TSP-1-dependent activation of TGF-β1 in prostates promoting the later stages of tumor growth, angiogenesis, and metastasis.

TSP-1 expression in the primary tumor is associated with diminished tumor growth due to inhibition of angiogenesis. In the early stages of tumor growth, TSP-1 expressed by stromal cells inhibits angiogenesis and tumor growth [40]. Breast cancer-derived tumors in TSP-1 null mice were significantly larger and heavier at 60 and 90 days of growth [41]. Similarly, we demonstrate that CaP tumors are significantly larger in TSP-1 null mice after only 12 days of growth. In the primary tumor, TSP-1 regulates angiogenesis, decreasing neovascularization and is upregulated by the tumor suppressors p53 and PTEN and inhibited by the oncogenes Myc and Ras [42]. The RM1 murine CaP cell line utilized in our studies was generated by overexpression of Myc and Ras; however this does not alter the cell line’s ability to express TSP-1 or induce angiogenesis. Our data demonstrate increased angiogenesis and blood vessel maturation in mice with TSP-1 deleted and this enhanced angiogenesis resulted in higher numbers of platelets within tumors. When TGF-β1 was deleted in platelets, angiogenesis was increased in tumors. These subcutaneous tumors had significantly higher numbers of mature, SMA covered blood vessels consistent with prior studies showing that platelet-deletion of TGF-β1 results in larger, more mature blood vessels [27]. Changes in tumor angiogenesis and maturation of blood vessels alter the numbers of platelets recruited into tumors.

Within tumors, platelets can stimulate migration and invasiveness of cancer cells. Our data demonstrate that the deletion of TSP-1 significantly increases the numbers of platelets recruited into tumors with a similar trend occurring after TGF-β1 deletion in platelets. This represents one method by which platelets could acquire tumor-derived proteins capable of controlling pre-metastatic niche formation and stimulating cancer cell migration into the circulation. In previous studies, we demonstrated that platelets regulated tumor-induced bone turnover and preferentially sequestered tumor-derived proteins [4, 5]. In this study, we show that platelets from TSP-1 null mice sequester TGF-β1 preventing its release and activation. Platelet-secreted TGF-β1 alters invasion and potentially hematogenous metastasis. High platelet counts were associated with higher levels of TGF-β1 in ovarian cancer patients and blockage of the TGF-β type I receptor prevented the platelet-induced invasion of ovarian cancer cells [43]. In breast cancer, TGF-β1 activation stimulates single-cell migration through the bloodstream, while a lack of TGF-β1 results in collective metastasis through lymphatic tissues in a tail vein injection model [44]. In another study, the treatment of breast cancer cells with platelet releasates induced a gene signature associated with epithelial-mesenchymal transition and enhanced metastatic potential, including MMP-9, VEGF, MCP-1, and tenascin, and activated TGF-β1 dependent pathways [11]. In colon cancer, platelet TSP-1 drives invasiveness through the induction of MMP-9 activity [45]. We further found that the MMP-9 concentration was higher in platelets after TSP-1 deletion or platelet-specific deletion of TGF-β1 similar to the induction of MMP-9 seen with platelet coculture with breast and colon cancer cells. However, the increase of MMP-9 in our study must results from other activation by other platelet-derived proteins as it occurs despite the absence of TSP-1 activation of TGF-β1.

Outside the primary tumor, TSP-1 and TGF-β1 are also found in platelets and serum, where they can interact with metastatic circulating tumor cells. Circulating TSP-1 enhanced metastasis in the presence of platelets [18] and increased TGF-β1 is associated with increased numbers of circulating tumor cells that drive metastasis [35]. In our study, we utilized the markers CXCR4 and CD117 to examine the numbers of circulating progenitor cells in the bloodstream. These two receptors are classic markers of cells residing in the bone marrow but are also postulated to be expressed on circulating tumor cells [30, 46]. Using a syngeneic model, we are unable to determine the origin of these cells definitively, but their numbers are indicative of pre-metastatic niche formation and progression of cancers toward a more aggressive phenotype which is being driven by platelet-sequestered proteins. Platelet TGF-β1 is necessary for breast cancer cell extravasation and colonization in the lungs [11]. Breast cancer cells adhere to TSP-1 and their growth and metastasis can be blocked with TSP-1 neutralizing antibodies [19]. TSP-1 stimulates metastatic cell colonization of the lung with the injection of TSP-1 doubling the metastatic rate. Platelets were partially required for this effect as depletion of platelets decreased the number of metastases even in the presence of exogenous TSP-1 [18]. In addition to activating latent TGF-β1 during platelet activation, TSP-1 also activates TGF-β1 in areas of shear stress [34]. Further, platelet-derived TSP-1 stimulates cancer cell adhesion [47]. These two processes represent one method by which the TSP-1/TGF-β1 axis might stimulate extravasation and colonization during the metastatic process. In our study, we were unable to measure metastatic cells. A majority of CaP cell lines do not spontaneously metastasize to bone and measurable metastases are unlikely to occur within the 12 days of tumor growth. Thus, we focus on the measurable pre-metastatic changes in the bone microenvironment induced by CaP growth.

In the bone microenvironment, the main metastatic site for CaP, TSP-1 and TGF-β1 are key signaling proteins controlling bone turnover. Even without tumors, we demonstrate that TSP-1 null control mice have a higher number of osteoclasts than WT control mice. Increased monocyte differentiation occurs in TSP-1 deficient mice [29]. The changes in monocyte differentiation likely correlate with changes in cells from the same lineage, namely macrophages and osteoclasts. When tumors were implanted, the numbers of osteoclast progenitors in the bone marrow were enhanced as demonstrated by increased *ex vivo* differentiation with MCSF and RANKL. In the bone marrow, TSP-1 is expressed on the endosteal surface, as well as in megakaryocytes and platelets [48]. On the other side of bone remodeling, TSP-1 induction of activated TGF-β1 induces osteoblast differentiation [49], representing one potential for the tumor-induced bone formation measured in the control animals. TGF-β1 stimulates early osteoblast proliferation while inhibiting osteoblast activation and bone formation and pushing mesenchymal stem cells towards chondrogenesis or myogenesis over osteogenesis [49]. Thus, alterations of theTSP-1/TGF-β1 signaling axis may regulate bone formation during osteoblastic versus osteoclastic bone lesions in CaP bone metastases.

TSP-1 and TGF-β1 both play integral roles in the interaction between metastatic CaP cells and the bone microenvironment. The co-culture of CaP cells with osteoblasts results in a significant increase in TSP-1 expression [50, 51], while co-culture with prostate stromal cells led to the downregulation of TSP-1 expression [51]. TGF-β1 regulation of bone cell differentiation and interaction with metastatic cancer cells is well documented. However, the role of TGF-β1 in the initial steps leading to metastasis is controversial and may be depended on the concentration of TGF-β1 and whether bone metastatic lesions are osteoblastic or osteoclastic. In breast cancer, neutralizing TGF-β1 prior to tumor implantation reduced the number and size of bone lesions and the associated osteolysis due to alterations in myeloid progenitor differentiation into osteoclasts [52, 53]. Neutralization of TGF-β1 resulted in decreased osteoclast number and function in both control and tumor-bearing animals which is opposite to the findings of our study. However, this neutralization would be more similar to a total knockout of TGF-β1 while our data is the result of TGF-β1 deletion in megakaryocytes and platelets alone leaving the expression of TGF-β1 by other bone microenvironmental cells. Conversely, silencing TGF-β1 in CaP cells reduced the incidence of osteoblastic lesions [54]. Our data demonstrate that TGF-β1 loss in myeloid-derived megakaryocytes and the platelets increase osteoclastogenesis and tipping the balance away from bone formation which may decrease the number of osteoblastic lesions. Supporting the role of the TSP-1 TGF-β1 axis in CaP-induce osteoclastogenesis, TGF-β1 treatment of LNCaP-C4-2B human CaP cells induced RANKL production likely stimulating osteoclast differentiation and function [55]. Further, MMP-9 expression by PC3 human CaP cells induces osteoclast recruitment and activation in the bone microenvironment [36]. Our data demonstrate increased platelet concentrations of both MMP-9 and RANKL in mice bearing CaP tumors after either total TSP-1 deletion or platelet-specific deletion of TGF-β1. Increases in either of these two proteins may be the cause of the enhanced osteoclastogenesis in mice bearing CaP tumors. Thus, the effect of TSP-1 induction on TGF-β1 activation and bone metastasis is likely dependent on whether cancer induces osteoclastic or osteoblastic lesions.

The utility of TSP-1 as a biomarker in the circulation is also controversial. Serum TSP-1 concentration was one of five biomarkers demonstrated to predict metastatic CaP [56], while other studies demonstrated TSP-1 staining and serum levels decrease with CaP progression [57, 58]. Accordingly, circulating TSP-1 levels were higher in women with metastatic breast cancer compared to those with localized disease, while tumoral expression of TSP-1 was increased in patients with early breast cancer [20]. TSP-1 platelet concentration when combined with free PSA to total PSA ratio and serum Cathepsin D concentrations was able to diagnose CaP [59]. Our data demonstrates that TSP-1 in platelets is increased in tumor-bearing mice and this is associated with an increase in platelet concentrations of TGF-β1. Circulating TGF-β1 is increased in patients with bone metastases and correlates with increased PF4 and other platelet-derived protein levels [4, 23]. In patients with metastatic breast cancer, high circulating TGF-β1 was associated with poor prognosis and increased numbers of circulating tumor cells [35]. In CaP patients, high preoperative plasma levels of TGF-β1 correlated with decreased progression-free survival [25]. Plasma TGF-β1 levels are largely due to the release of platelet TGF-β1 during the collection and preparation of blood samples [27]. Clinically, whether platelet activation likely occurs prior to testing depends upon the anti-coagulant used and the time between collection and processing. For our studies, blood is collected into EDTA containing prostaglandin E_1_ to minimize platelet activation and separated with eight hours of collection for human samples and within two hours for murine samples. Thus, the concentration of the proteins within the platelets should be representative of the concentration circulating in platelets *in vivo*. Additional studies are needed to determine whether changing the collection procedures would alter the utility of platelet-derived TSP-1 or TGF-β1 as potential biomarkers for cancer progression.

In summary, we demonstrate that targeting the TSP-1/TGF-β1 axis may not alter primary tumor growth but could be used to block pre-metastatic communication between the primary tumor and bone microenvironment, possibly preventing metastasis. Reduction of TSP-1 or platelet-derived TGF-β1 stimulated osteoclastogenesis and reduced tumor-induced bone formation indicating a potential role for the TSP-1/TGF-β1 axis in the development of osteoblastic or mixed CaP lesions although examination of bone metastatic CaP would be required. The ability to target platelet-sequestered proteins could decrease the toxicities seen with systemic perturbation of either TSP-1 or TGF-β1. Finally, the presence and concentration of TSP-1 and/or TGF-β1 in platelets represent a potential biomarker for aggressive cancers although further studies would be required examining a wider patient population.

## MATERIALS AND METHODS

### Reagents

All chemicals were obtained from Sigma (St. Louis, MO) unless otherwise specified. Media were supplied by the Central Cell Services Core of the Lerner Research Institute.

### Cell Culture

RM1 murine prostate cancer cells (RRID: CVCL_B459) were provided by Dr. W.D. Heston (Cleveland Clinic). These cells were generated from C57BL/6 mice using the mouse prostate reconstitution model with overexpression of *Myc* and *Ras* [60, 61]. RM1 tumors provide insight on aggressive, androgen-independent, and metastatic CaPs [62]. Cells were cultured in RPMI 1640 supplemented with 10% heat-inactivated FBS, 100 U/mL penicillin, 100 µg/mL streptomycin and were regularly passaged by trypsinization (0.05% trypsin, 0.53 mM EDTA).

### Tumor Cell Injection

Age-matched, male 8-12 week old C57BL/6J (WT; RRID: ISMR_JAX:000664), B6.129S2-*Thbs1^tm1Hyn^*/J (TSP-1 null; RRID: ISMR_JAX:006141) (Jackson Laboratory, Bar Harbor, ME), PF4-Cre^-/-^ TGFβ1^fl/fl^ (TGFβ1^fl/fl^), or PF4-Cre^+/-^ TGFβ1^fl/fl^ (PF4-Cre^+^ TGFβ1^fl/fl^, a kind gift from Dr. Barry Coller), mice were anesthetized by 100 mg/kg ketamine and 10 mg/kg xylazine injected i.p.. RM1 cells at a concentration of 4 x 10^5^ cells in 0.1 mL PBS were injected s.c. into the flank of mice on each side. Control mice of each background were injected s.c. in each flank with 0.1 mL PBS. Tumors were allowed to grow 12 days before sacrifice. Animal experiments were performed under Cleveland Clinic IACUC approved protocols (2011-0655; 2015-1369). Mice were maintained on standard chow in standard housing.

### Plasma and Platelet Isolation

While mice were anesthetized, the vena cava was exposed, and blood was collected from the vein into an acid-citrate-dextrose buffer containing 1 µg/mL prostaglandin E_1_ (Sigma, St. Louis, MO). Platelets and plasma were separated from the platelet-rich plasma of blood by gel filtration, as previously described [6]. The activated supernatant was collected by centrifugation after platelets were treated with 100 nM phorbol 12-myristate 13-acetate (PMA) (Sigma), 1 mM protease inhibitors (Roche complete Mini, Indianapolis, IN), and 500 µM mouse PAR-4-amide (Bachem, Torrance, CA) at 37°C for 45 minutes to stimulate release of granular contents.

### Immunoblotting

Activated platelet releasates and platelet lysates were incubated with 5X Lane Marker Reducing sample buffer (Pierce) and boiled at 95°C for 10 min. Samples underwent electrophoresis on a 7% polyacrylamide gel at 100 V for approximately 2 hours. Proteins were transferred to a nitrocellulose membrane at 100 V for 1 hour at 4°C. Membranes were blocked in 10% milk in TBS containing 0.1 % Tween-20. TSP-1 antibody (Santa Cruz Biotechnology; RRID: AB_831756) was used to detect platelet-derived proteins, with fibrinogen antibody (Abcam; RRID: AB_732366) used as a loading control. Horseradish-peroxidase conjugated secondary antibodies (BioRad; RRIDs: AB_11125345 and AB_808614) were used with a luminescent substrate (Pierce SuperSignal West Pico) to visualize proteins.

### Tumor Analysis

Tumors were removed from the mice upon sacrifice. Tumors were weighed, and the dimensions of the tumors were measured by caliper. Tumor volume was calculated as (d_1_ x d_2_)^(3/2) x (π/6), as described by Voeks et al. [62]. Tumor density was calculated as tumor weight/tumor volume. Tumors were then placed in 10% formalin or embedded in OCT for sectioning. Tumors embedded in paraffin were sectioned and stained with hematoxylin and eosin. Tumors embedded in OCT freezing medium were sectioned at 7µm. Sections were then fixed in 4% PFA and incubated with primary antibodies: anti-CD31 (1:50, BD Pharmigen; RRID: AB_393571), anti-CD42b (1:75, Millipore; RRID: AB_10712312) or anti-smooth muscle actin (SMA, 1:100, Abcam; RRID: AB_2223021). Tissues were then washed in PBS and exposed to a fluorescently labeled secondary antibody (Invitrogen). Slides were mounted with Vectamount+DAPI (Vector Labs). Slides were scanned with an Olympus VS110, and whole tumor staining analyzed using the Halo Software.

### Lymphocyte Isolation

Lymphocytes were isolated from whole blood using Ficoll-Paque PLUS (GE Healthcare, Piscataway, NJ) according to the manufacturer’s procedure. Briefly, whole blood was layered on Ficoll-Paque PLUS and centrifuged at 2000 rpm for 10 minutes without brakes. The lymphocyte layer was removed and washed twice in 10 mL 1X PBS by centrifugation at 1200 rpm. Pelleted lymphocytes were resuspended in 2 mL DMEM/F12 and stored at 4°C until staining.

### Flow Cytometry

Isolated lymphocytes were blocked with 2 µL murine FcR blocking reagent (Pierce) for 15 minutes. Lymphocytes were then incubated with 20µL of fluorescently conjugated antibodies. Either CD184/CXCR4 conjugated with FITC (R&D Systems; RRID: AB_2091799) or CD117/c-kit conjugated with PE (Miltenyi Biotec; RRID: AB_2660118) were used. Stained lymphocytes were then washed twice with 1X PBS. Lymphocytes were then resuspended in 500 µL 1% formalin. Stained lymphocytes were then analyzed using a BD FACS Canto II running the FACS Diva software (RRID: SCR_001456). Fluorescence values of stained lymphocytes were normalized to an isotype control sample.

### Microcomputed Tomography (MicroCT)

The microCT analysis was performed one day prior to the injection of tumor cells and after 12 days of tumor growth. Pre-injection scans were performed in live animals with *ex vivo* scans performed at experimental termination on isolated legs in PBS after overnight fixation in 10% formalin. Scans were conducted in the Cleveland Clinic Biomedical Imaging and Analysis Core Center on a GE eXplore Locus microCT (GE Healthcare, Piscataway, NJ) and 360 X-ray projections were collected in 1° increments (80 kVp; 500 mA; 26 min total scan time). Projection images were preprocessed and reconstructed into 3-dimensional volumes (1024^3^ voxels, 20 µm resolution) on a 4PC reconstruction cluster using a modified tent-FDK cone-beam algorithm (GE reconstruction software). Three-dimensional data was processed and rendered (isosurface/maximum intensity projections) using MicroView (Parallax Innovations). For each volume, a plane perpendicular to the z-axis/tibial shaft was generated and placed at the base of the growth plate. A second, parallel plane was defined 1.0 mm below, and the entire volume was cropped to this volume of interest for quantitative analysis [63]. Image stacks from each volume of interest were exported for quantitative analysis. Cancellous bone masks were generated in MicroView, and 3D trabecular structural indices were extracted using custom MatLab (The MathWorks, Inc, Natick, MA; RRID: SCR_001622) algorithms. Trabecular thickness (Tb.Th) and trabecular spacing (Tb.Sp) were determined by previously reported methods. The trabecular number (Tb.N) was calculated by taking the inverse of the average distance between the medial axes of trabecular bone segments. Bone volume fraction (BV/TV, total bone voxels divided by total cancellous bone mask voxels) and bone surface area (BSA, the sum of pixels along edges of trabecular bone) were also calculated for each VOI.

### ELISAs

Plasma and activated platelet releasates were subjected to Blue Gene Bone Alkaline Phosphatase, R&D Systems TRANCE/RANKL, or R&D Systems MMP-9 ELISA assays to measure the concentration of proteins. A StanBio Total Calcium LiquiColor Procedure No. 0150 Assay was used to measure calcium levels in the plasma samples.

### Bone Histology

To visualize changes in the bone structure, we isolated the tibiae from mice after microCT scanning. Tibiae were isolated from whole legs, fixed in 10% formalin, and decalcified in 14% neutral buffered EDTA for 10d. Bones were then cut longitudinally and washed with 1X PBS. Bones were then dehydrated in serially increasing concentrations of ethanol. Dehydrated bones were cleared in xylene and embedded in paraffin. Paraffin-embedded bones were sectioned at 5 µm and placed on slides. Sections were stained with hematoxylin and eosin or for 30 minutes in the TRAP solution described below. Slides were scanned with the Hamamatsu NanoZoomer by the Virtual Microscopy Core in the Wake Forest School of Medicine. Bone histomorphometry, osteoclast numbers, and growth plate organization were analyzed with the BioQuant Osteo software (RRID: SCR_016423).

### Bone Marrow Isolation and Osteoclast Differentiation

Tibiae were isolated from mice, and the ends were removed. The bone marrow (BM) was extruded with PBS containing penicillin and streptomycin. The BM cell suspension was filtered through a cell strainer and washed. Isolated BM cells were plated at a density of 1×10^6^ cells in 6 well plates in αMEM media containing 10% FBS, penicillin and streptomycin, and 50 ng/mL MCSF. Seven days later, media was replaced with complete αMEM treated with 25 ng/mL murine MCSF and 30 ng/mL RANKL (R&D Systems) to induce osteoclastogenesis. Media with treatments were changed every few days for 2-3 wks.

### TRAP Staining

Tartrate-resistant acid phosphatase (TRAP) staining was performed to determine the number of multinucleated osteoclasts formed from isolated bone marrow macrophage cells. Cells were washed and fixed in a 10% glutaraldehyde solution for 15 min at 37°C. Cells were then incubated with a TRAP staining solution (0.3 mg/mL Fast Red Violet LB, 0.05 M sodium acetate, 0.03 M sodium tartrate, 0.05 M acetic acid, 0.1 mg/mL naphthol, 0.1% Triton X-100, pH 5.0) for 2 hours at 37°C. Cells were observed using a Leica DMIRB inverted light microscope in the LRI Imaging Core. TRAP-stained cells with at least three nuclei were counted per well and were considered differentiated osteoclasts.

### Statistical Analysis

Student’s *t* test analysis or one-way ANOVA analysis with Tukey post-test were used to determine statistical significance using the GraphPad Prism 7.0 software (La Jolla, CA; RRID: SCR_002798). * represents p<0.05, ** represents p<0.01, *** represents p<0.005.

## Abbreviations

CaP: prostate cancer
MMP: matrix metalloproteinase
RANKL: receptor activator of NF-κB ligand
SMA: smooth muscle actin
TGF-β1: transforming growth factor beta-1
TRAP: tartrate-resistant acid phosphatase
TSP: thrombospondin
VEGF: vascular endothelial growth factor

## ONLINE SUPPLEMENTAL MATERIAL

Supplemental methods, references, figures, and legends.

## AUTHOR CONTRIBUTIONS

Investigation: BAK, KSH, LS; Writing-Original Draft: BAK; Writing-Review & Editing: BAK, KSH, LS, JSW, DSP, TVB; Resources: JSW, DSP; Formal analysis and Visualization: BAK; Conceptualization, Funding acquisition, Project administration, Supervision: BAK, TVB

## ACKNOWLEDGMENT

We thank Miroslava Tischenko, Richard Rozic, and Steven Maximuk for technical assistance. Dr. Barry Coller of Rockefeller University provided the PF4-Cre/TGFβ1 floxed mice. Dr. Amit Vasanji developed the software and techniques for microCT analysis.

---

**Supplementary Figure 1.**
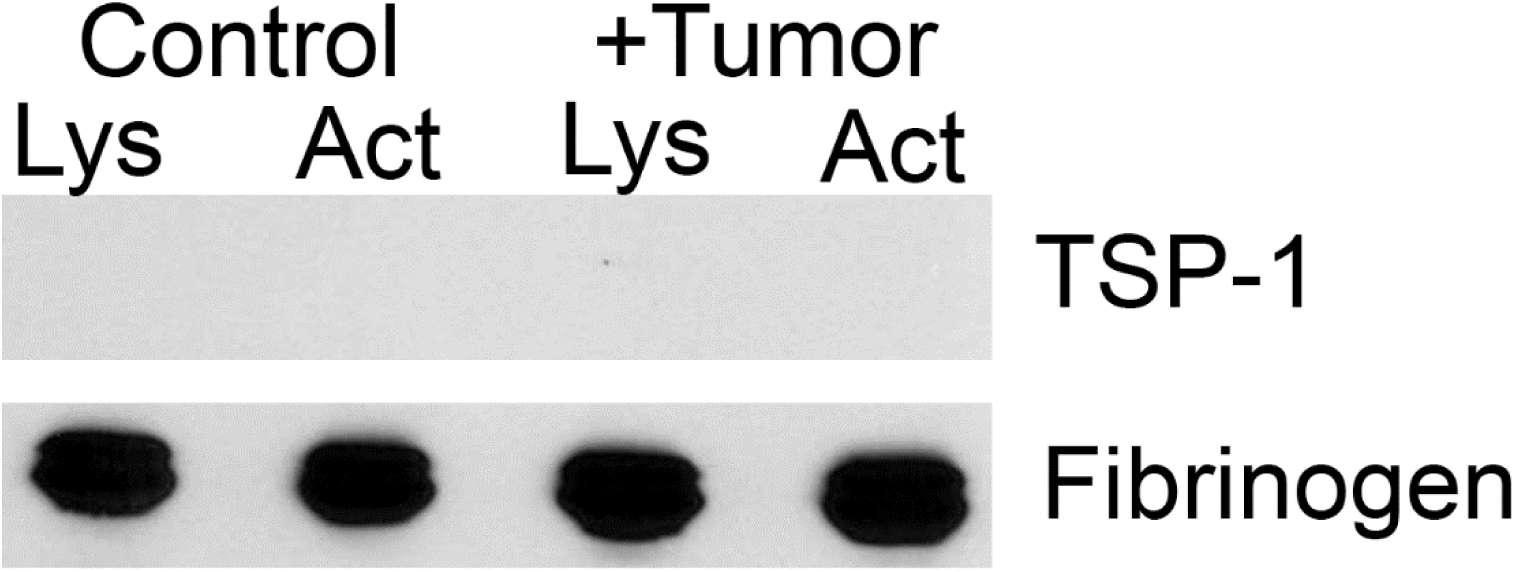

**Supplementary Figure 2.**
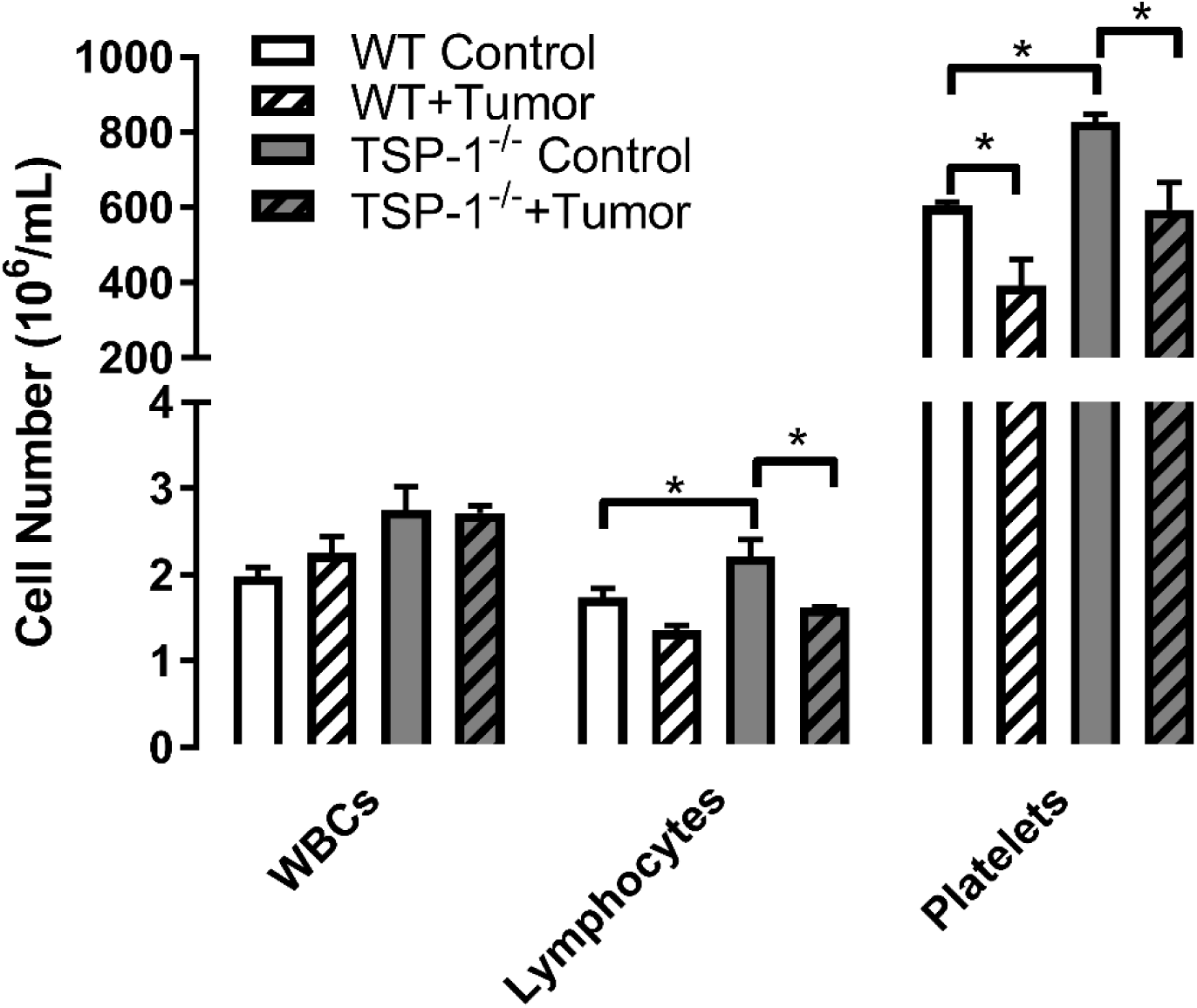

**Supplementary Figure 3.**
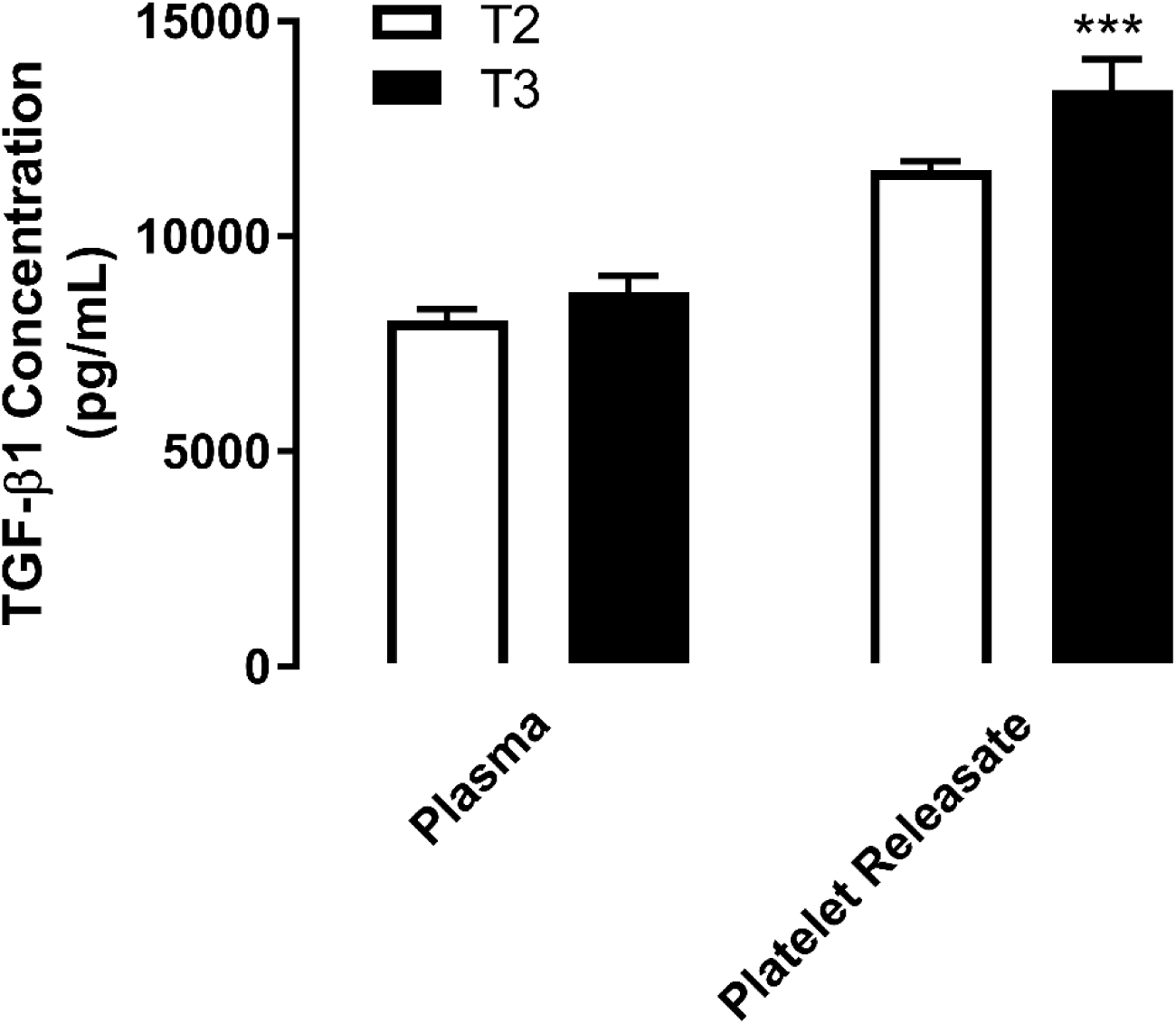

