## Supplementary Material for "Platelet TSP-1 Controls Prostate Cancer-Induced Osteoclast Differentiation and Bone Marrow-Derived Cell Mobilization through TGFβ-1"

**SUPPLMENTARY MATERIAL**

SUPPLEMENTAL METHODS

*Whole Blood Analysis*

Mice were anesthetized with 10 mg/kg xylazine and 100 mg/kg ketamine. The vena cava of mice was exposed and blood was collected from the vein into 0.1 mL acid-citrate-dextrose (ACD) buffer containing 1 µg/mL $PGE_1$. Whole blood was subjected to complete blood count (CBC) analysis with differential to determine blood cell concentrations using an Advia 120 laser hematology system (Siemens Healthcare Diagnostics) in the Center for Cardiovascular Research at the Lerner Research Institute. White blood cell (WBC) counts, lymphocyte counts, platelet counts, and mean platelet volume (MPV) were assessed.

*Clinical Samples*

An approval from the Cleveland Clinic Institutional Review Board was obtained prior to the initiation of blood sample collection from patients undergoing radical prostatectomy at the Cleveland Clinic Glickman Urological and Kidney Institute. A patient consent form was specifically created for the study with clearly stated goals and describing the purpose of our research. Whole blood (3-4 ml) was collected by venipuncture in $Na_2EDTA$ tubes (BD Biosciences, San Jose, CA) from patients prior to surgery. Plasma and platelets were isolated from whole blood by centrifugation and gel filtration, as previously described (Kerr et al., 2010, 2013). Platelets were activated with 50 µM human TRAP-6-amide (Bachem), 1 mM protease inhibitors, and 100 nM PMA to stimulate granule release. Tumor sections from each patient were analyzed by a pathologist and T stages were assigned.

*ELISAs*

Isolated plasma and platelet releasates from six patients were assayed using the RayBiotech Human TGF-β1 ELISA according to the manufacturer's instructions. Two separate samples from each patient at each time point were analyzed and compared to a standard curve to obtain the concentration in pg/mL of each protein in the samples.

SUPPLEMENTARY FIGURE 1

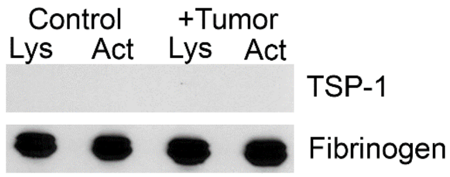


**Supplementary Figure 1: Platelets derived from TSP-1 null mice are TSP-1 deficient.** TSP-1 null mice were injected subcutaneously with RM1 cells or mock injected with PBS. Platelets were isolated from the whole blood of mice after 12 days. Platelets were lysed in RIPA/PI or activated with PMA and PAR-4 to stimulate release of granular contents. Activated releasates and platelet lysates were analyzed for TSP-1 protein expression by immunoblotting. Fibrinogen was used as a loading control.

SUPPLEMENTARY FIGURE 2

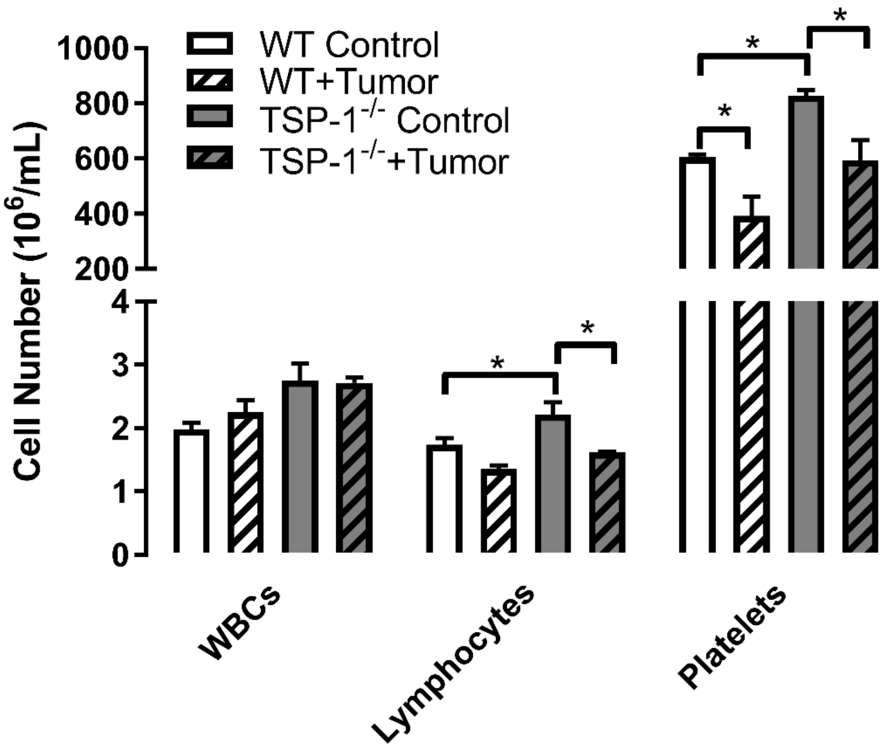


**Supplementary Figure 2: Whole blood analysis of TSP-1 null and WT mice in the presence and absence of RM1 tumors.** TSP-1 null and WT mice were injected subcutaneously with RM1 cells or mock injected with PBS (Control). Whole blood was isolated after 12 days. White blood cell (WBC), lymphocytes, and platelet cell numbers were measured and represented as mean cell number±SEM ($n$=3). * represents $p<0.05$ by one-way ANOVA.

SUPPLEMENTARY FIGURE 3

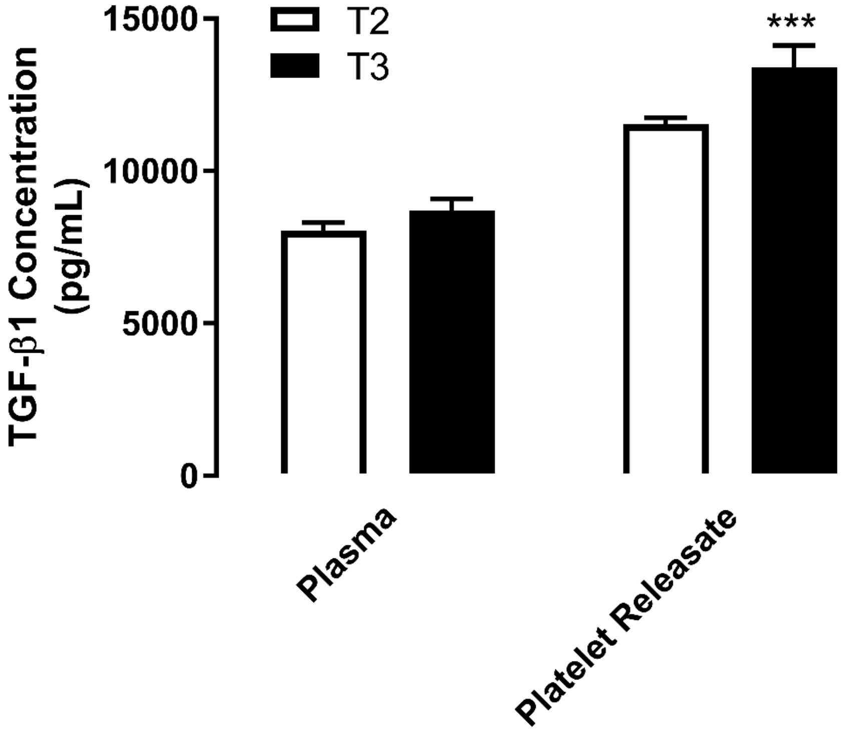


**Supplementary Figure 3: Plasma and platelet TGF-β1 concentration in low and high staged prostate cancer patients.** Plasma and releasate from platelets activated with 50 μM TRAP-6 and 100 nM PMA were isolated from whole blood collected from patients undergoing radical prostatectomy and mean TGF-β1 concentration±SEM of combined plasma and releasate concentrations analyzed by ELISA are shown ($n$=3). T2 represents samples from low stage, less aggressive prostate cancer patients and T3 represents samples from high stage, aggressive prostate cancer patients as determined by a pathologist. *** represents $p<0.005$ by Student's $t$ test.
